# Contributions of the Heme Coordinating Ligands of the *Pseudomonas aeruginosa* Outer Membrane Receptor HasR to Extracellular Heme Sensing and Transport

**DOI:** 10.1101/2020.04.26.062984

**Authors:** Alecia T. Dent, Angela Wilks

**Author notes:** To whom correspondence should be addressed: Angela Wilks, Department of Pharmaceutical Sciences, School of Pharmacy, University of Maryland, HSF II, 20 Penn Street, Baltimore, MD 21201-1140.

## Abstract

*Pseudomonas aeruginosa* exhibits a high requirement for iron which it can acquire via several mechanisms including the acquisition and utilization of heme. *P. aeruginosa* encodes two heme uptake systems, the <u>h</u>eme <u>a</u>ssimilation <u>s</u>ystem (Has) and the *<u>P</u>seudomonas* <u>h</u>eme <u>u</u>tilization (Phu) system. Extracellular heme is sensed via the Has system that encodes an extra cytoplasmic function (ECF) σ factor system. Previous studies have shown release of heme from the extracellular hemophore HasAp to the outer membrane receptor HasR is required for activation of the σ factor HasI. Herein, employing site-directed mutagenesis, allelic exchange, quantitative PCR analyses, immunoblotting and ^13^C-heme uptake studies, we characterize the differential contributions of the outer membrane receptor HasR extracellular FRAP/PNPNL loop residue His-624 and the N-terminal plug residue His-221 to heme transport and signaling, respectively. Specifically, we show mutation of the N-terminal plug His-221 disrupts both signaling and transport. The data is consistent with a model where heme release from HasAp to the N-terminal plug of HasR is required to initiate signaling, whereas His624 is required for simultaneously closing off the heme transport channel from the extracellular medium and triggering heme transport. Furthermore, mutation of His-221 leads to dysregulation of both the Has and Phu uptake systems suggesting a possible functional link that is coordinated through the ECF σ factor system.

## INTRODUCTION

Iron is an essential micronutrient that is required for the survival and virulence of almost all bacterial pathogens. Iron is extremely limited in mammalian hosts due to sequestration by the innate immune system through upregulation of iron storage proteins, down regulation of ferritin, and the secretion of iron chelating proteins such as lipocalin-2 (1). During infection, bacterial pathogens including *Pseudomonas aeruginosa* circumvent iron deficiency by utilizing a variety of iron acquisition systems (2-4). In response to iron starvation, *P. aeruginosa* deploys two siderophore systems, pyoverdine (Pvd) and pyochelin (Pch) (5, 6) as well as the ferrous uptake system (Feo) (7). *P. aeruginosa* also encodes two non-redundant heme acquisition systems, the *Pseudomonas* heme uptake (Phu) and heme assimilation (Has) systems that utilize heme as an iron source (8, 9).

Bacteria respond and adapt to extracellular stimuli through cell-surface signaling (CSS) systems which are coupled to extra cytoplasmic function (ECF) σ factors (10, 11). These alternative σ factors complex with the core RNA polymerase, direct binding to the promoter of a target gene, and activate transcription. *P. aeruginosa* encodes several iron-responsive ECF σ factors including PvdS, FpvI and HasI (12-14). We have recently shown that the Has system which encodes the ECF σ factor HasI is primarily required for sensing extracellular heme whereas the Phu system is the major heme transporter (9). The Has system senses heme through the secreted extra cellular hemophore HasAp which on binding to the TonB-dependent OM receptor HasR triggers the CSS. Recent studies employing site-directed mutants of HasAp revealed the activation of the CSS requires heme release from HasAp and capture by HasR (15). Capture of the heme by HasR transduces the signal through the N-terminal plug of HasR resulting in inactivation of the anti-σ factor HasS, release of the ECF σ factor HasI, and transcriptional activation of the *hasRADEF* operon. As extracellular heme levels decrease, HasS rapidly sequesters HasI down regulating the system.

Once heme is transported across the outer membrane by either HasR or PhuR it is translocated to the cytoplasm by the PhuT-PhuUV periplasmic ABC transport system. Within the cytoplasm, heme is specifically trafficked to the iron-dependent heme oxygenase (HemO) by the cytoplasmic heme binding protein PhuS. HemO catalyzes the oxidative cleavage of heme to release CO, iron and the heme metabolites biliverdin IXβ (BVIXβ) and - IXδ (BVIXδ) (17). In addition to heme-dependent transcriptional activation of the Has system, the *hasAp* transcript is subject to post-transcriptional regulation by the heme metabolites BVIXβ and/or BVIXδ (16).

Heme uptake pathways associated with ECF σ factor systems are conserved across a number of bacterial pathogens including *Serratia marcescens* (18,19), *Bordetella pertussis* (20), and *P. aeruginosa* (8). The significance of heme sensing and uptake in *P. aeruginosa* pathogenesis is evident in the fact that clinical isolates from patients with chronic lung infection have adapted to utilize heme while decreasing pyoverdine biosynthesis (21). The Has system has several layers of transcriptional and post-transcriptional regulation that allows the bacteria to rapidly fine tune its response to changes in the host environment (15,16). In an attempt to dissect the mechanism of heme signaling and transport by the *P. aeruginosa* Has system, we have employed site directed mutagenesis, allelic exchange, qPCR, immunoblotting and ^13^C-heme uptake studies to determine the contributions of the heme coordinating ligands of HasR (Fig. 1). Previous studies employing similar approaches with variants of HasAp showed heme release from HasAp is triggered on simultaneous release of the Tyr-75 and His-32 heme ligands (15).

**Figure 1.**
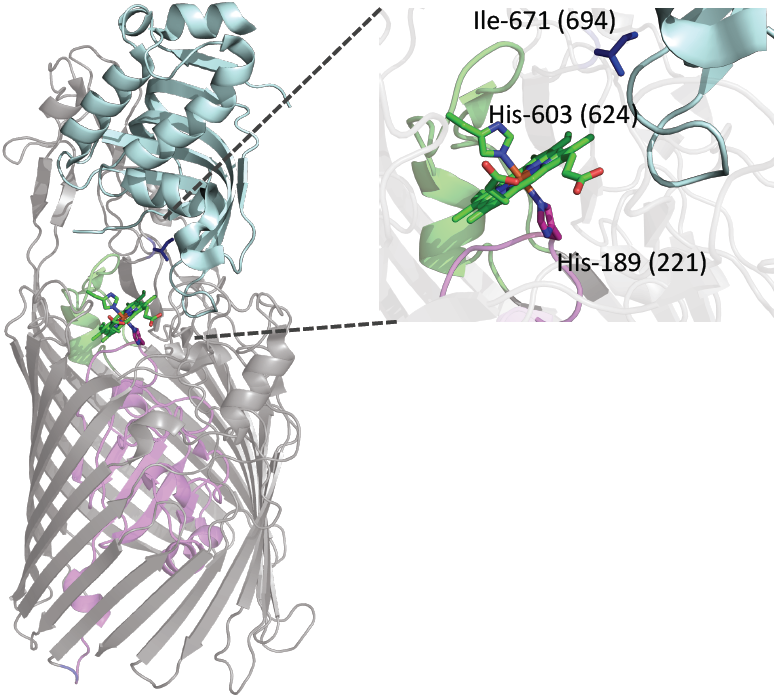
Structure of holo-HasAp-HasR complex showing HasAp (light blue) and HasR with the N-terminal plug (magenta) and β-barrel (grey) (1). The heme (green) and heme-coordinating residues His-189 (magenta), His-603 on FRAP/PNPNL loop (green) and Ile-671 (dark blue) are shown as sticks. The number in parentheses represent the corresponding residues in *P. aeruginosa* HasR. Image generated from PDB file 3CSL in PyMOL (29).

Herein we show the heme coordinating residue (His-624) of the conserved FRAP/PNPNL loop and the N-terminal plug heme coordinating residue (His-221) are required for heme transport (Fig 1). The FRAP/PNPNL loop so called for the conserved amino acid motifs is proposed to close off the extracellular milieu from the heme transport channel. Similarly, Ile-694 located on extracellular loop L8, previously proposed to prevent heme backsliding to HasAs, is also essential for heme transport (Fig 1). In contrast the N-terminal plug His-221 is critical for both heme signaling and transport. Furthermore, heme bound to HasAp is not accessible to PhuR as shown by the significant growth defect in the transport deficient HasR mutants when supplemented with holo-HasAp rather than heme. Interestingly, the N-terminal plug *hasRH221R* strain, which is unable to either activate the CSS cascade or transport heme, revealed a global dysregulation in both the Has and Phu system indicating a direct or indirect link between the ECF σ factor HasI and the Phu heme uptake system.

## RESULTS

### The hasR allelic strains show similar HasR and HasAp expression profiles in low iron

The expression profiles of the *hasR* allelic strains were first analyzed in iron deplete conditions. The *hasR* variant strains all show a similar growth profile to PAO1 WT (Fig 2A). For the *hasR* loop mutant strains, the *hasR* mRNA levels at each timepoint were all within 2-fold of the parent PAO1 strain, with negligible differences in protein expression (Fig 3A and B; Fig S1A). The data confirms the HasR loop variants are folded and stable. Similarly, the *hasAp* mRNA and protein levels for the *hasR* loop variants were very similar to those in the PAO1 WT strain (Fig 3C and D; Fig S1B). In contrast to the *hasR* loop variants, the N-terminal plug *hasRH221R* strain showed a significant increase in HasR protein at the later timepoints (Fig 3B and S1A). To confirm that the expression of the Phu system was not compromised in the *hasR* variant strains we analyzed the mRNA and protein levels by qPCR and Western blot, respectively. The loop *hasRH624A, H624Y*, and *I694G* variants showed similar mRNA and protein profiles as the PAO1 WT strain (Fig 4). Interestingly, the relative mRNA levels of *phuR* in the *hasRH221R* strain revealed a significant increase over PAO1 WT (Fig 4A). Furthermore, this increase in relative mRNA levels translated to protein expression (Fig 4B). Overall, the data suggests that the *hasRH221R* variant even in low iron conditions has a distinct phenotype from that of the *hasR* loop variants.

**Figure 2.**
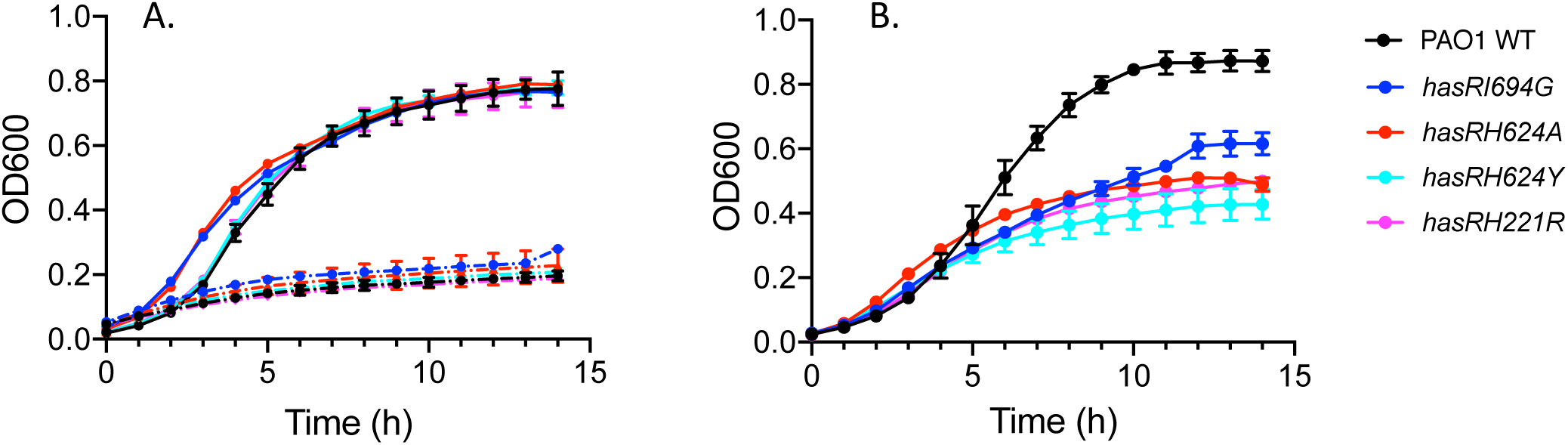
Growth curves of PAO1 WT and the *hasR* allelic strains. *A.* Strains were grown in M9 minimal media (dashed line) or M9 supplemented with 1 μM heme (solid line). *B.* Cells grown in M9 minimal media supplemented with 1 μM holo-HasAp. Strains are color coded as shown in the legend.

**Figure 3.**
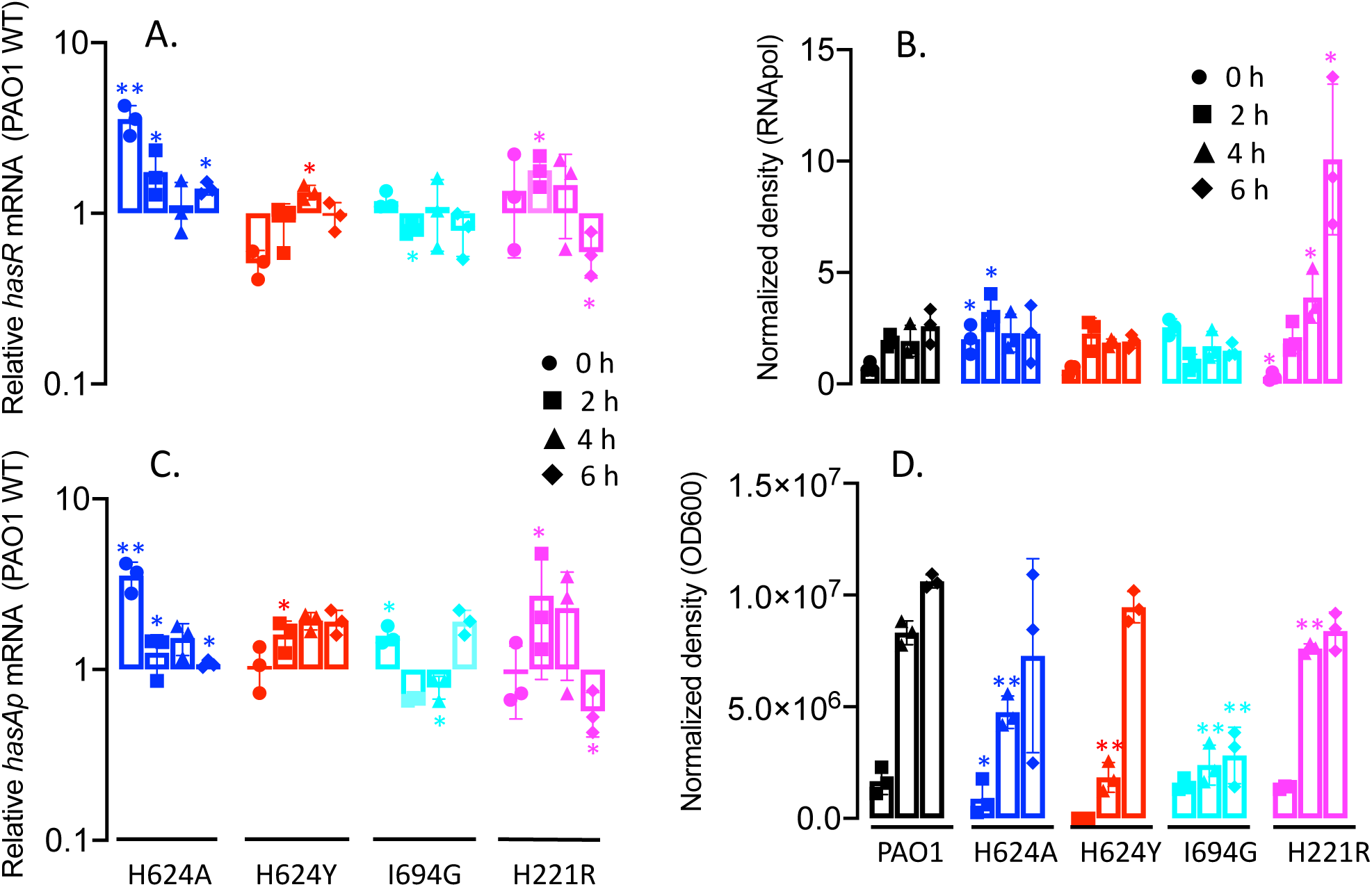
Relative HasR and HasAp mRNA and protein levels for the *hasR* allelic strains in iron-deplete conditions. *A. hasR* mRNA isolated at 0, 2, 4, and 6 h following growth in M9 minimal media. mRNA values represent the mean from three biological experiments each performed in triplicate and normalized to PAO1 WT at the same timepoint. *Error bars* represent the standard deviation from three independent experiments performed in triplicate. The indicated *p* values were normalized to mRNA levels of PAO1 WT at the same timepoint, where *, *p* < 0.05. *B.* Western blot analysis of PAO1 WT and the *hasR* allelic strains. For HasR total protein (25 μg) was loaded in each well. RNA polymerase α was used as a loading control. Normalized density (*n* = 3) was performed for three separate biological replicates. The indicated *p* values were normalized to PAO1 at the same timepoint where *, *p* <0.05, **, *p* < 0.005. *C. hasAp* mRNA analyzed as in A. *D.* Western blot analysis of HasAp as in B. Extracellular supernatant (4 μL) was loaded and run on the automated Wes capillary western system as described in the Experimental Procedures. Representative Western blot images are shown in the Supporting Information (Fig S1).

**Figure 4.**
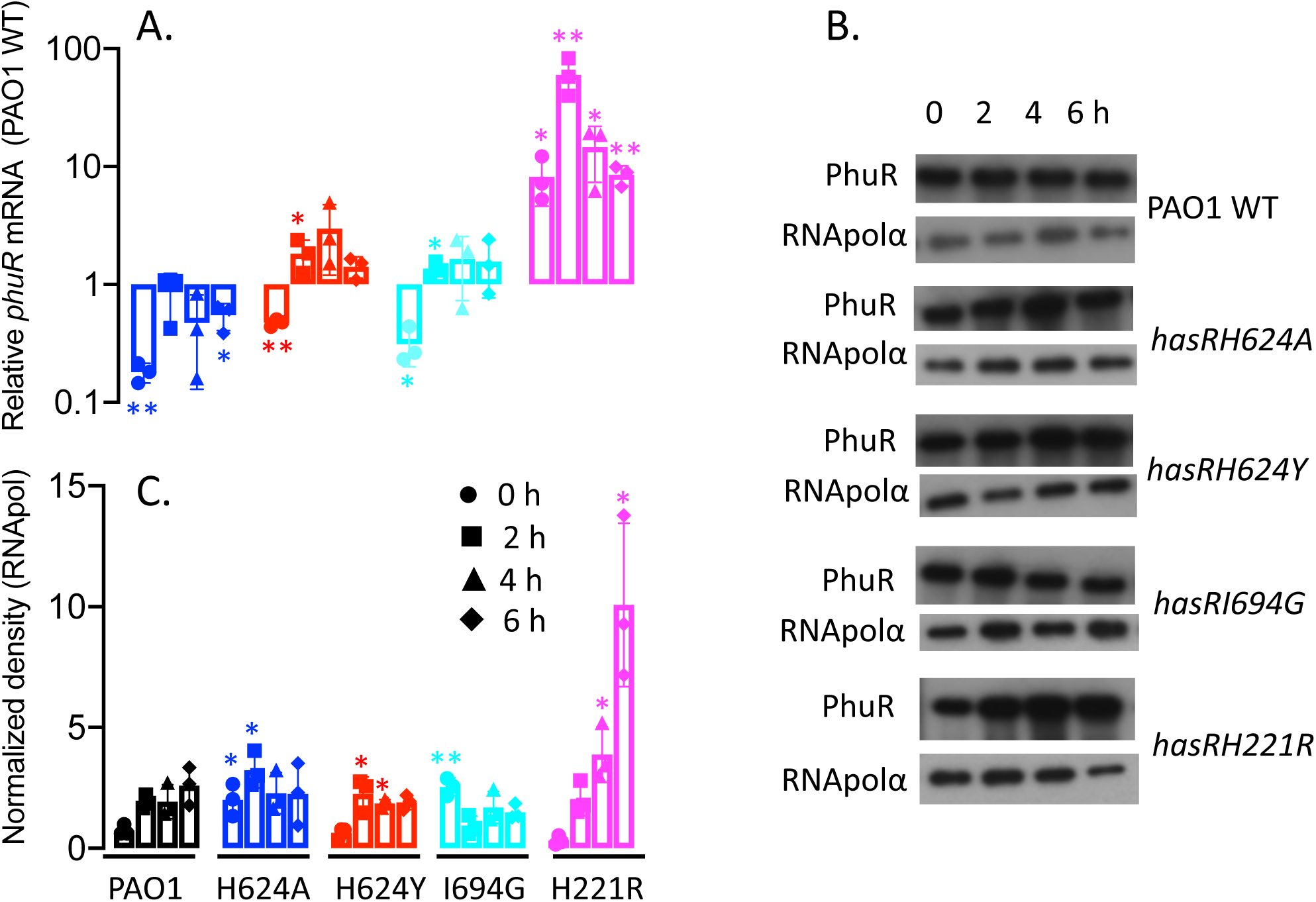
Relative *phuR* mRNA and protein levels for the *hasR* allelic strains in iron-deplete conditions. *A*. mRNA isolated at 0, 2, 4, and 6 h following growth in M9 minimal media. mRNA values represent the mean from three biological experiments each performed in triplicate and normalized to PAO1 WT at the same timepoint. *Error bars* represent the standard deviation from three independent experiments performed in triplicate. The indicated *p* values were normalized to mRNA levels of PAO1 WT at the same timepoint, where *, *p* < 0.05; **, *p* < 0.005. *B.* Representative Western blot of PAO1 WT and the *hasR* allelic strains. Total protein (25 μg) was loaded in each well. RNApolα was used as a loading control. *C.* Normalized density (*n* = 3) was performed on Western blots for three separate biological replicates. The indicated *p* values were normalized to PAO1 WT at the same timepoint where *, *p* <0.05; **, *p* < 0.005.

### The hasR allelic strains are deficient in heme uptake

The growth of the *hasRH624A, H624Y, I694G* and *hasRH221R* strains were monitored following supplementation with either 1 μM heme or 1 μM holo-HasAp. All of the mutant strains were competent to utilize “free” heme as an iron source when compared to the growth rates in iron-deplete media (Fig 2A). However, when heme was supplied in the form of holo-HasAp a significant inhibition in growth was observed for all of the variant strains compared to PAO1 WT (Fig 2B). These results suggest that heme but not heme complexed to HasAp is accessible to the *hasR* variant strains via the Phu heme uptake system. qPCR analysis of *phuR* on heme supplementation show the relative mRNA levels of the *hasR* loop variants are similar to PAO1 WT at all timepoints (Fig S2). However, the relative mRNA levels in the *hasRH221R* variant are ten-fold higher than those of PAO1 at all timepoints, consistent with the profile in iron-deplete conditions. The PhuR protein levels for PAO1 WT and all of the *hasR* variant strains follow a similar profile where the protein is induced at 2 h, followed by a decrease at 4 h and a subsequent increase again at 6 h. This pattern of expression we interpret as a shock response on addition of free heme, as we do not observe this effect when heme is complexed to HasAp. However, despite statistically significant differences in protein expression levels as determined by Western blot, it is clear that all of the *hasR* variant strains express PhuR protein within 2-fold of that observed for PAO1 WT (Fig S2B and C).

The ability of the *hasR* variant strains to utilize heme, but not holo-HasAp, was confirmed by isotopically labeled ^13^C-heme uptake and ICP-MS experiments. Supernatants were collected 4 h following supplementation of PAO1 WT or the *hasR* allelic strains with either ^13^C-heme or ^13^C-heme-HasAp. The resulting BVIX isomers were extracted and quantified by LC-MS/MS. When supplemented with ^13^C-heme the *hasR* variant strains were all capable of utilizing heme as judged by the similar levels of ^13^C-BVIXβ and ^13^C-BVIXδ compared to PAO1 WT (Fig 5A). Consistent with the ability to utilize exogenously added ^13^C-heme as an iron source, ICP-MS analysis confirmed the intracellular iron content for all of the *hasR* variant strains is similar to PAO1 WT (Fig 5B). Taken together the data suggests that in the absence of a functional Has system “free” heme is readily acquired by the major transporter PhuR.

**Figure 5.**
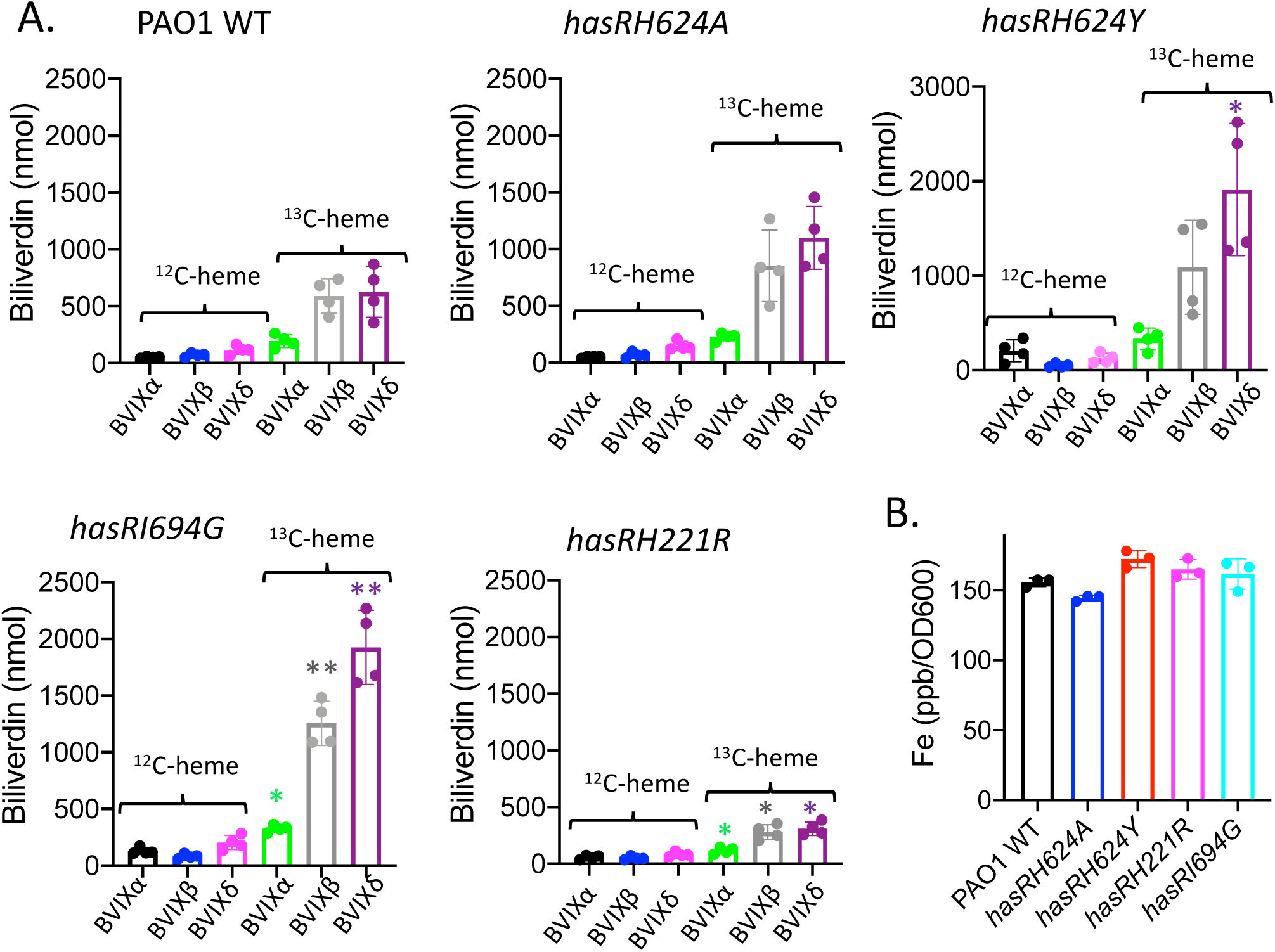
Heme utilization by the *hasR* allelic strains when supplemented with ^13^C-heme. *A.* LC-MS/MS analysis of the BVIX isomers supplemented with 1 μM ^13^C-heme at 4 h. Biliverdin values represent the standard deviation of four biological replicates. The indicated *p* values for each strain compared back to PAO1 WT for the respective BVIX isomers where *, *p* < 0.05; **, *p* < 0.005. *B.* ICP-MS analysis of intracellular iron levels. The indicated *p* values for each strain compared back to PAO1 WT where *, *p* < 0.05; **, *p* < 0.005.

In contrast to supplementation with ^13^C-heme, the *hasR* variant strains were unable to utilize the ^13^C-heme-HasAp complex as an iron source (Fig 6A). Concomitant with the decreased ability to utilize extracellular heme, we see a slight but significant increase in endogenous heme (^12^C-heme) metabolism. Such redistribution of endogenous heme iron is consistent with the slower growth rate and dysregulation of iron homeostasis (Fig 2B). The inability to utilize heme bound to HasAp was further confirmed on measurement of the intracellular iron levels by ICP-MS (Fig 6B). The *hasR* variant strains all showed decreased intracellular iron content, which is particularly noticeable for the H221R N-terminal plug mutant. Analysis of *phuR* mRNA levels in the *hasR* loop variants are within 2-fold of those for PAO1 WT (Fig. S3). Western blot further confirmed protein levels mirror those of PAO1 (Fig S3B and C). However, as can be seen from both the qPCR and Western blot analysis the relative mRNA and protein levels are significantly higher for the *hasRH221R* plug variant, particularly at the 6 h timepoint (Fig S3). This difference is not solely due to the decreased iron levels (Fig 6B), as iron depletion is also observed for the *hasR* loop variants. Furthermore, we observed a similar mRNA and protein expression profile for the *hasR* variant strains in cultures supplemented with heme, where the intracellular iron levels were identical to PAO1 WT (Fig 5B). Taken together the data clearly shows heme bound to HasAp, in the absence of a functional HasR transporter, is not accessible to PhuR. This is consistent with previous studies in which we have shown the Has and Phu systems have distinct non-redundant roles in heme signaling and transport, respectively (9).

**Figure 6.**
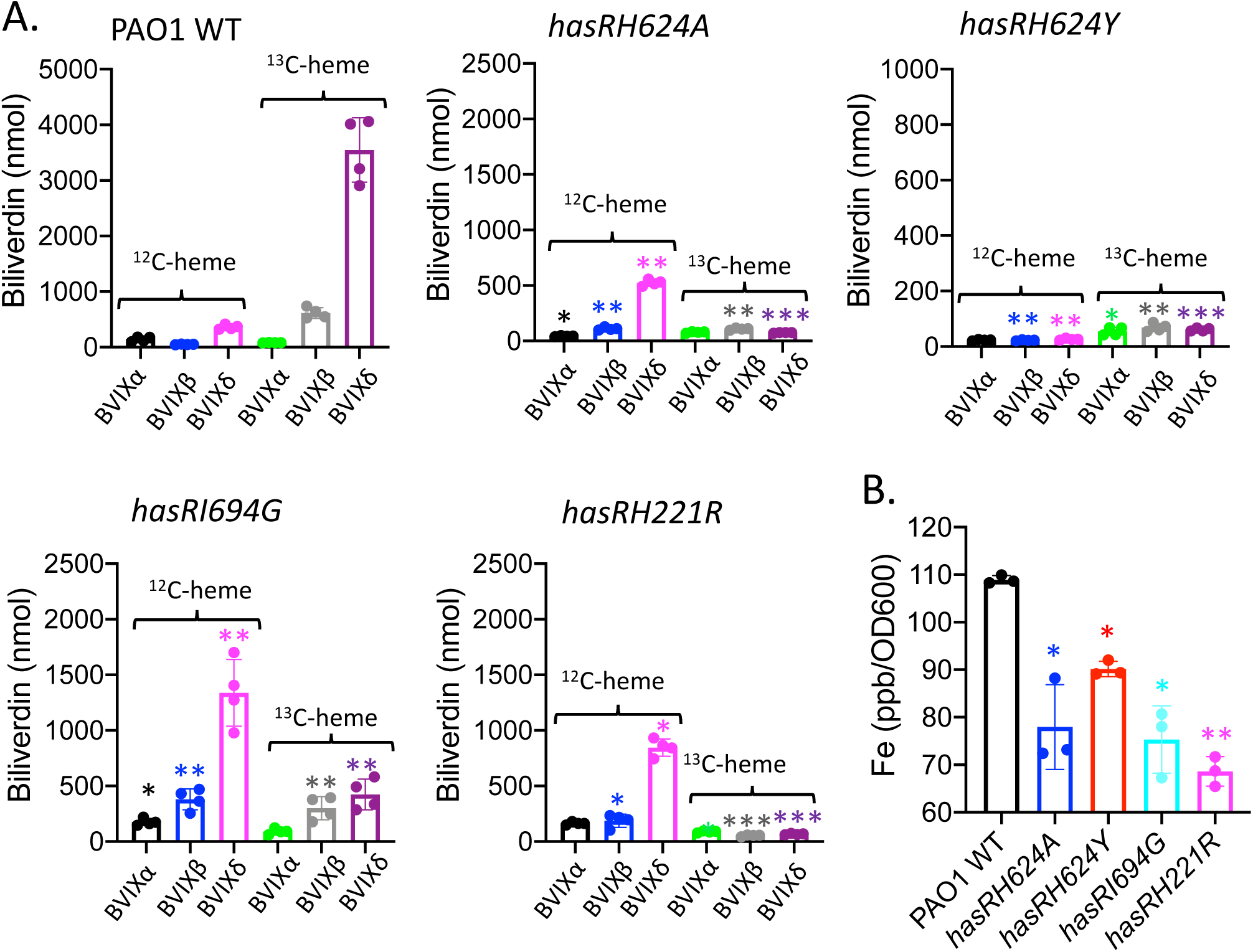
Heme utilization by the *hasR* allelic strains when supplemented with ^13^C-heme-HasAp. *A.* LC-MS/MS analysis of the BVIX isomers supplemented with 1 μM ^13^C-heme-HasAp at 4 h. Biliverdin values represent the standard deviation of four biological replicates. The indicated *p* values for each strain compared back to PAO1 WT for the respective BVIX isomers where *, *p* < 0.05; **, *p* < 0.005.*B.* ICP-MS analysis of intracellular iron levels. The indicated *p* values for each strain compared back to PAO1 WT where *, *p* < 0.05; **, *p* < 0.005.

### Activation of ECF σ factor system by the hasR H624A, H624Y, I694G and H221R allelic strains

We next sought to determine the contributions of the extracellular loop and N-terminal plug residues on activation of the CSS cascade. Following supplementation with either heme or holo-HasAp, HasI dependent transcriptional activation of *hasR* and *hasAp* was monitored over time by qPCR analysis and Western blot. As previously reported supplementation of PAO1 WT with 1 µM heme after an initial decrease in mRNA levels we observed an increase in *hasR* mRNA levels over time as extracellular HasAp levels accumulate (Fig 7A). The *hasAp* mRNA profiles mirror those of *hasR*, however, as previously reported the relative mRNA levels are greater than *hasR* due to increased mRNA stability following processing of the *hasRAp* transcript (Fig 7B) (16). Western blot analysis of HasR and HasAp protein levels are consistent with the respective mRNA profiles (Fig 8).

**Figure 7.**
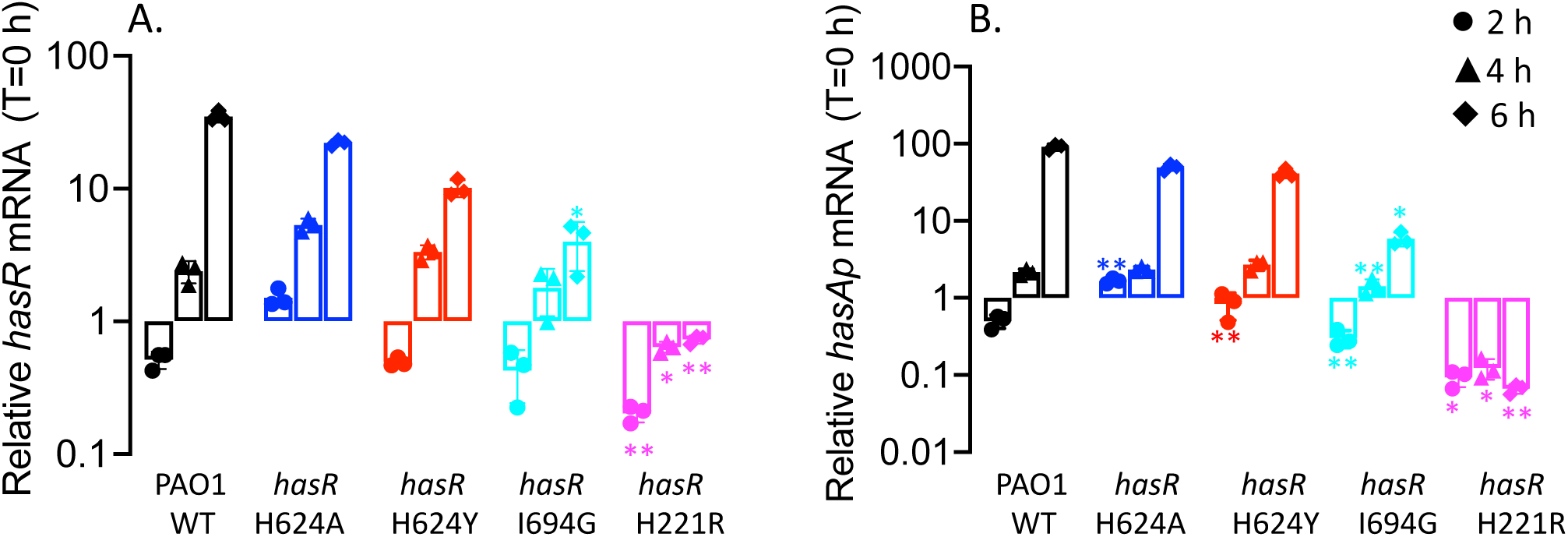
Relative *hasR* and *hasAp* mRNA levels for the *hasR* allelic strains in heme supplemented conditions. *A. hasR* relative mRNA levels. *B. hasAp* relative mRNA levels. mRNA isolated at 0, 2, 4, and 6 h following growth in M9 supplemented with 1μheme. mRNA values represent the mean from three biological experiments each performed in triplicate and normalized to PAO1 WT at the same timepoint. *Error bars* represent the standard deviation from three independent experiments performed in triplicate. The indicated *p* values were normalized to mRNA levels of PAO1 WT at the same timepoint, where *, *p* < 0.05; **, *p* < 0.005.

**Figure 8.**
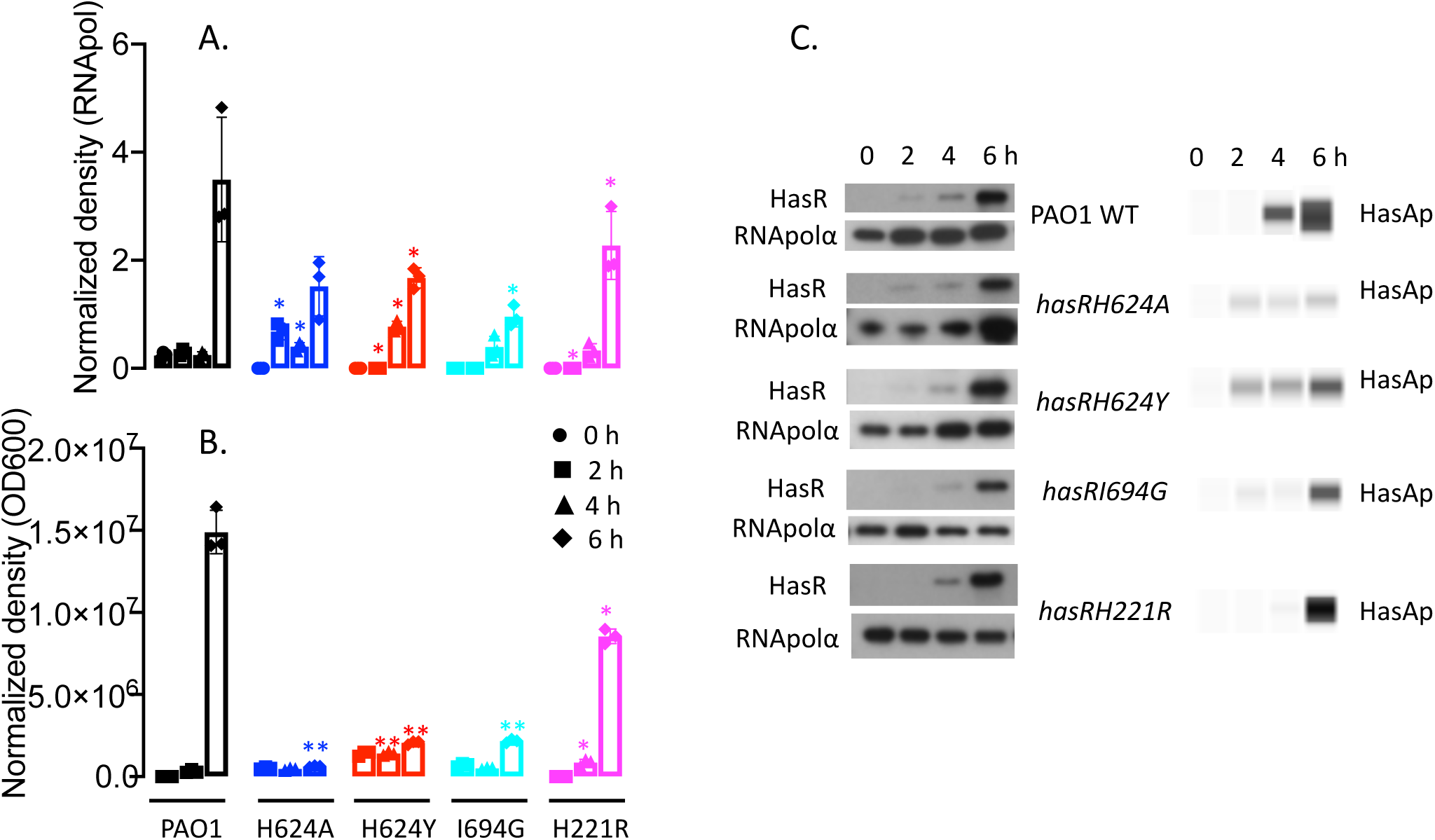
Relative HasR and HasAp protein levels for the *hasR* allelic strains in heme supplemented conditions. *A*. HasR protein at 0, 2, 4, and 6 h following growth in M9 minimal media supplemented with 1 μM heme. Total protein (25 μg) was loaded in each well. RNA polymerase α was used as a loading control. Normalized density (*n* = 3) was performed on Western blots for three separate biological replicates. *B.* HasAp as in A. Extracellular supernatant (4 μl) was loaded for analysis on the automated Wes capillary system as described in Experimental Procedures. Normalized density (*n* = 3) was corrected for differences in OD600 for three biological replicates. *Error bars* represent the standard deviation from three independent experiments performed in triplicate. Indicated *p* values for the *hasR* allelic strains normalized to PAO1 WT at the same timepoint, where *, *p* < 0.05; **, *p* < 0.005. *C.* Representative Western blot of PAO1 WT and the *hasR* allelic strains for HasR and digital HasAp images generated by capillary western analysis on an automated Wes system.

The *hasR* loop variants show a similar profile but lower *hasR* and *hasAp* mRNA and protein levels compared to WT (Fig 7 and 8). The increase in relative mRNA levels on addition of heme suggests that despite an inability to transport heme, they retain the ability to activate the CSS cascade (Fig 7). In contrast the *hasRH221R* strain showed no heme-dependent transcriptional activation of *hasR* and *hasAp* (Fig 7). This lack of transcriptional activation is not solely the result of decreased HasAp levels in the extracellular medium, as we observe significant protein levels by Western blot (Fig 8). Previous studies from our laboratory have shown *hasAp* is subject to positive post-transcriptional regulation by the heme metabolites BVIXβ and/or BVIXδ (15, 16). Therefore, the discrepancy between mRNA and protein can be rationalized by the fact heme actively taken up by the Phu system is further metabolized to BVIXβ and BVIXδ.

We further analyzed the transcriptional activation of the *has* operon on supplementation with a fixed concentration of holo-HasAp. In these studies, we utilized a previously characterized functional but truncated HasAp to distinguish, by Wes analysis, the supplemented holo-HasAp from the endogenously produced protein (15). In contrast to the lag on addition of heme (∼4 h; see Fig 7), addition of holo-HasAp to PAO1 immediately activates transcription from the *hasR* promoter (Fig 9A). For PAO1 WT, transcriptional activation peaks at the earlier 2 h timepoint, and is maintained over 6 h. In contrast the *hasR* loop variants show decreased transcriptional activation at the early timepoints. The lag in transcriptional activation for the loop variants is further manifested at the protein level, where significantly higher HasR levels are seen at the earlier timepoints for PAO1 WT (Fig 10A). Whereas the *hasAp* mRNA levels for the *hasR* loop variants follows a similar trend to those of *hasR* (Fig 9B), we detect barely any endogenous HasAp protein (Fig 10B). This lack of protein expression can be rationalized by the inability of the *hasR* loop variants to uptake heme leading to lower intracellular BVIX levels and a decrease in HasA translational efficiency. For the N-terminal plug *hasRH221R* strain we observe no transcriptional activation of the *hasR* operon, as shown by the lack of increase in *hasR* mRNA over time (Fig 9A). As expected we also observe no transcriptional activation of *hasAp* at the 2 h timepoint (Fig 9B). However, the accumulation of *hasAp* mRNA at the later timepoints is a consequence of both the inherent stability of the *hasAp* mRNA transcript versus *hasR*, and derepression of Fur as a consequence of iron deficiency (Fig 6B). Indeed, the relative increase in *hasR* and *hasAp* mRNA levels over time is also observed for the *hasR* loop variants, suggesting iron deficiency is also contributing in some degree to the increase in transcription. Furthermore, this translates to HasR expression where we observe an increase in protein levels at the 4 and 6 h timepoints (Fig 10A). Particularly noticeable in the *hasRH221R* strain are the detectable levels of HasAp despite the fact we see no accumulation of BVIX metabolites (Fig 10B). Although this could be attributed to the lower intracellular iron levels the lack of detectable HasAp protein in the *hasR* loop variants (that are also iron deficient) does not support this (Fig 6B). Furthermore, the increase in PhuR protein expression in the *hasRH221R* strain over both the WT and *hasR* loop variants in all conditions, is more consistent with a global dysregulation in iron homeostasis.

**Figure 9.**
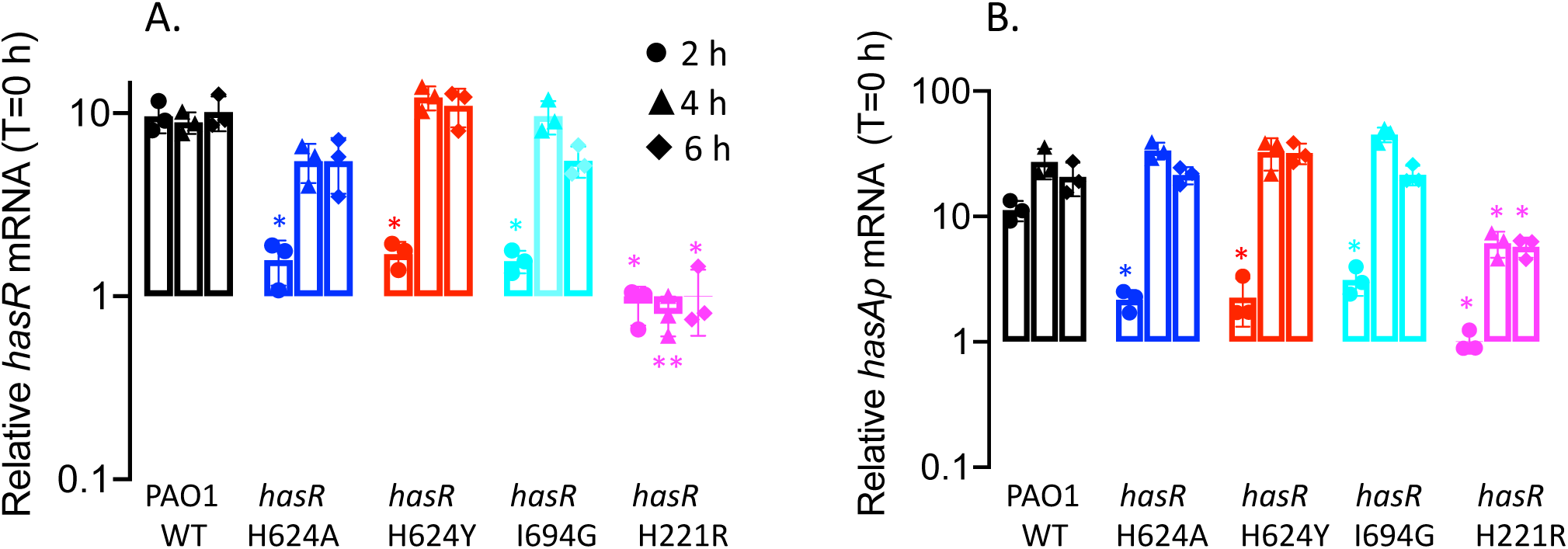
Relative *hasR* and *hasAp* mRNA levels for the *hasR* allelic strains in holo-HasAp supplemented conditions. *A. hasR* relative mRNA levels. *B. hasAp* relative mRNA levels. mRNA isolated at 0, 2, 4, and 6 h following growth in M9 supplemented with 1 μM holo-HasAp. mRNA values represent the mean from three biological experiments each performed in triplicate and normalized to time 0 h for each strain. *Error bars* represent the standard deviation from three independent experiments performed in triplicate. The indicated *p* values were normalized to mRNA levels of PAO1 WT at the same timepoint, where *, *p* < 0.05; **, *p* < 0.005.

**Figure 10.**
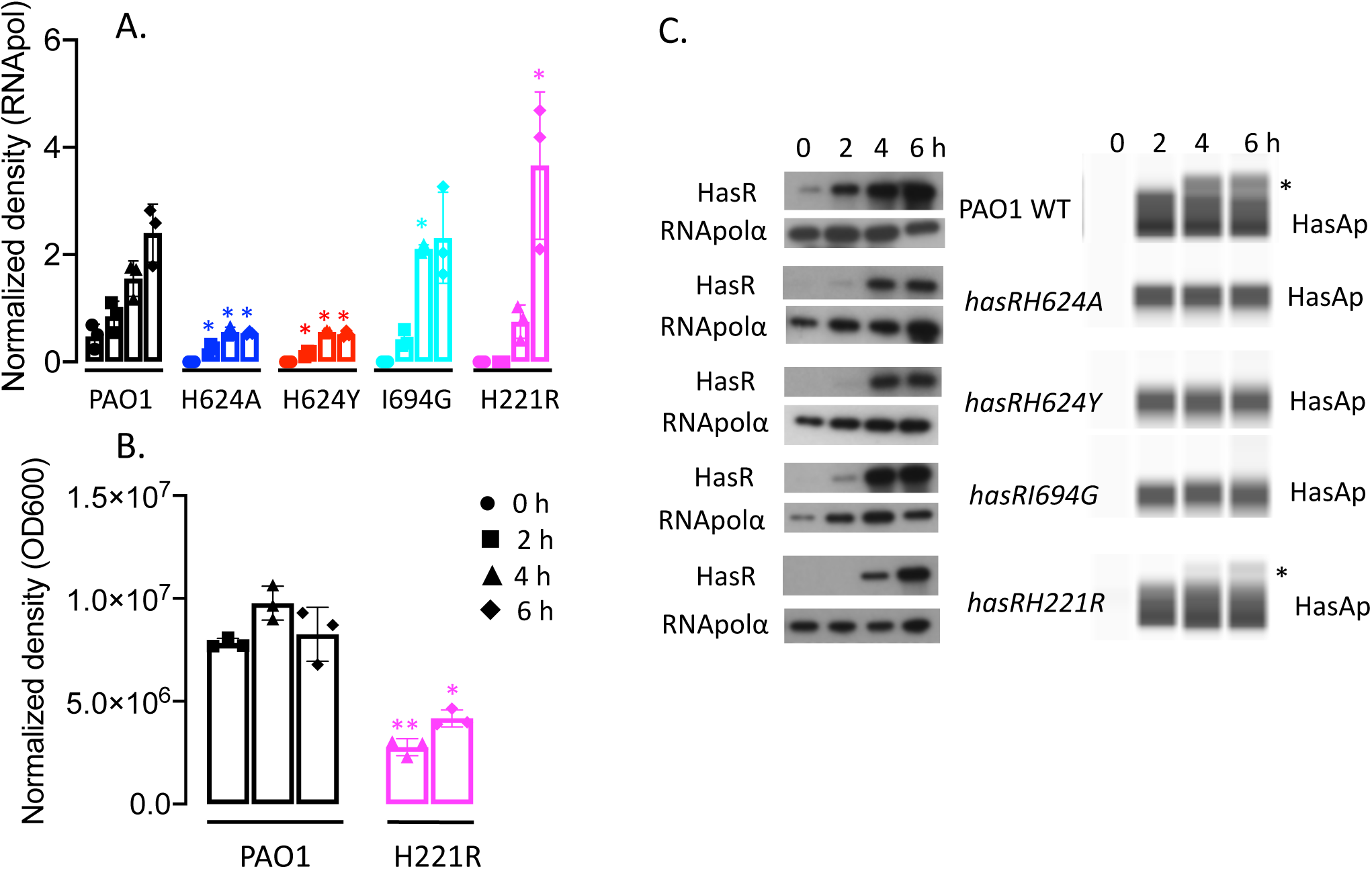
Relative HasR and HasAp protein levels for the *hasR* allelic strains in holo-HasAp supplemented conditions. *A*. HasR protein at 0, 2, 4, and 6 h following growth in M9 minimal media supplemented with 1 μM holo-HasAp. *B.* As in A for HasAp in the extracellular supernatant. *Error bars* represent the standard deviation from three independent experiments performed in triplicate. The indicated *p* values for HasR were normalized to RNA polymerase α subunit as a loading control, and to OD600 for HasAp. Indicated *p* values for the *hasR* allelic strains normalized to PAO1 WT at the same timepoint, where *, *p* < 0.05; **, *p* < 0.005. *C.* Representative Western blot of PAO1 WT and the *hasR* allelic strains for HasR and HasAp digital images generated by capillary western analysis on an automated Wes system. Normalized density (*n* = 3) was performed on Western blots for three separate biological replicates. The indicated *p* values were normalized to PAO1 WT at the same timepoint where *, *p* <0.05; \*\**p* < 0.005.

## DISCUSSION

The crystal structure of the *S. marcescens* HasAs-HasR complex revealed His-603 of the extracellular FRAP/PNPNL loop (L7) and His-189 on the N-terminal plug as the heme coordinating ligands (Fig 1) (22). Based on the crystal structure, HasAs was proposed to transfer heme through a transient intermediate where His-32 of HasAs is displaced on protein-protein interaction with HasR, while the stronger Tyr-75 ligand remains coordinated due to deprotonation of Tyr-75 by the nearby His-83 (22). In subsequent steps, steric hindrance by Ile-671 of HasR, and transient protonation by His-83 promotes release of heme to HasR. It was also suggested that Ile-671 sterically hinders heme from backsliding into the HasAs hydrophobic pocket (Fig 1). However, our recent studies with the *P. aeruginosa* Δ*hasAp* strain supplemented with exogenous holo-HasAp H32A, Y75A or H83A mutant proteins showed an increase in the transcriptional activation of the CSS cascade compared to the WT strain (15). The heme content in the holo-HasAp WT and mutant proteins was identical, suggesting that the increased transcriptional activation in the mutant proteins resulted from a kinetically trapped intermediate. This data suggested that in contrast to the sequential release of His-32 and subsequent protonation of Tyr-75, heme release from HasAp occurs via a concerted mechanism, where both His-32 and Tyr-75 are simultaneously released through conformational rearrangement and free energy gain on interaction with HasR (15). To further study the mechanism of heme transfer, uptake and initiation of the Has CSS cascade we turned our attention to the role of the OM receptor HasR in heme signaling and transport. Specifically, the role of the heme ligands His-221 of the N-terminal plug, His-624 of the FRAP/PNPNL loop, as well as the extracellular loop (L8) Ile-694 proposed to promote heme release from HasAp.

The *hasRH624A* strain on supplementation with holo-HasAp was unable to utilize heme judged by lack of ^13^C-heme metabolites and decreased intracellular iron levels (Fig 6), but still able to trigger the activation of the CSS cascade (Fig 9). The data indicates that heme coordination by the N-terminal plug His-221 is sufficient to trigger the CSS cascade, although significantly less efficiently than the WT strain. The data further suggests that heme coordination to HasR via the FRAP/PNPNL loop His-624 and N-terminal plug His-221 is required to trigger conformational rearrangement of the plug to facilitate heme transport, while simultaneously closing off the receptor channel from the extracellular environment.

To further test the role of heme coordination on heme signaling and transport we mutated the FRAP/PNPNL loop His-624 to a Tyr mimicking the heme coordination motif of PhuR (23). Similarly, we observed the *hasRH624Y* strain on supplementation with holo-HasAp was capable of triggering the CSS cascade but deficient in heme uptake. Given that heme complexed to HasAp is not accessible to the PhuR receptor (Fig 2 and 6) one might have expected making the receptor more “PhuR-like” would disrupt heme signaling and transport. However, the fact we observe activation of the CSS cascade (Fig 9) shows heme can be displaced from holo-HasAp, suggesting the inability of PhuR to extract heme from HasAp is more likely due to lack of a direct protein-protein interaction. However, somewhat surprisingly we observed no active heme uptake indicating that either Tyr-624 is i) sterically hindered from coordinating to the heme and allowing loop closure, or ii) the Tyr-His coordination is not conducive to triggering the conformational rearrangement of the N-terminal plug required for transport. Biochemical and spectroscopic characterization of the purified HasR variants reconstituted in lipid nanodiscs is currently being performed to determine their heme coordination properties.

Interestingly, despite the fact that the FRAP/PNPNL heme coordinating residue His-624 and the N-terminal plug His-221 remain intact, the *hasRI694G* strain when supplemented with holo-HasAp is unable to transport heme (Fig 6). Similar to the His-624 mutants, qPCR analysis revealed the *hasRI694G* strain was still capable of activating the ECF σ factor HasI, albeit with an apparent decrease in efficiency, as judged by the lower relative mRNA levels compared to WT (Fig 7 and 9). Taken together the data is somewhat consistent with the proposed model where the extracellular loop (L8) Ile-671 promotes steric displacement of heme from HasAs to HasR, while also preventing heme from backsliding to HasAs (22). Our current data on the *P. aeruginosa* HasR further suggests that loss of Ile-694 still allows for transient release of heme from HasAp to the receptor, such that heme signaling is triggered, but heme is not sterically constrained to allow loop closure and transport. Taken together the data shows both His-624 and Ile-694 are essential for facilitating heme release, capture and transport. Ongoing biochemical and spectroscopic characterization of the purified HasR variants reconstituted in lipid nanodiscs will shed more light on the effect of the FRAP/PNPNL loop mutations on heme binding and coordination.

In contrast to the *hasR* loop variant strains the *hasRH221R* variant strain was deficient in both heme signaling (Fig 7 and 9) and transport (Fig 6). It is unlikely that holo-HasAp does not interact with the HasR H221R receptor, given that we observe release of heme to the *hasR* loop variants which requires physical interaction with HasAp. Therefore, heme coordination to His-221 is absolutely required for transduction of the signal from the periplasmic face of the N-terminal plug to the cytoplasmic signaling domain. Taken together the data is consistent with a model where conformational rearrangement on interaction of holo-HasAp with HasR, facilitated by the steric clash of Ile-694 triggers heme release to HasR. Following release, heme binding to the HasR N-terminal plug via His-221 triggers the signal through the N-terminal domain, inactivating the anti-σ factor HasS, releasing the σ factor HasI, and transcriptionally activating the *has* operon. Simultaneous release of HasAp and loop closure activates heme transport through the receptor channel to the periplasmic face.

Interestingly, we initially attempted to construct the *hasRH221A* variant strain, however, on sequencing we noted an inversion of the DNA sequence within this region that led to a significant change in the amino acid sequence (data not shown). This suggested the *hasRH221A* mutation introduced structural changes within the plug that most likely resulted in the strain not being viable. Therefore, we constructed the *hasRH221R* strain incorporating the more conservative His to Arg substitution. These observations suggest that subtle conformational changes within the N-terminal plug may potentially lead to destabilization of the receptor or uncoupling of the anti-σ factor HasS. Interestingly, even in the *hasRH221R* strain we observe a distinct phenotype from that of the *hasR* extracellular loop variant strains, including significant differences in the expression profile of PhuR. As previously mentioned in iron deplete conditions the relative mRNA and protein levels of PhuR are significantly higher than either the WT or *hasR* extracellular loop variants (Fig 3). Furthermore, on closer analysis of the PhuR Western blots when supplemented with heme or holo-HasAp a doublet can be observed in what appears to be post-transcriptional or proteolytic processing (Fig S2 and S3). Taken together the data suggests that inactivation of the CSS cascade, and the cells ability to sense extracellular heme, has a direct or indirect effect on the Phu system. Although the Has and Phu systems have distinct non-redundant roles in heme sensing and transport, it is reasonable to assume that there is a regulatory link given their distinct roles in adapting to changes in physiological environments. It is unclear at this time the nature of the regulatory link between the Has and Phu systems but it is possible that the ECF σ factor HasI has as yet undetermined targets beyond auto regulation of the *has* operon.

Furthermore, the mechanistic insight into heme release, signaling and transport gained from these studies provides a platform for future structural and mechanistic studies of the heme transport channels of both HasR and PhuR.

## EXPERIMENTAL PROCEDURES

### Bacterial strains, plasmids and growth conditions

Bacterial strains and plasmids used in this study are listed in Table S1 and oligonucleotide primers and probes in Table S2. *E. coli* strains were routinely grown in Luria Bertani (LB) broth (American Bioanalytical) or on LB agar plates, and *P. aeruginosa* strains were freshly streaked and maintained on *P.* isolation agar (PIA) (BD Biosciences). Brain heart infusion agar and LB were used in generating the *hasR* mutant strains as described previously (57). All strains were stored frozen at −80 °C in LB broth with 20% glycerol. For qPCR and Western blotting, singly isolated colonies from each *Pseudomonas* strain were picked, inoculated into 10 mL of LB broth, and grown overnight at 37 °C with shaking (210 rpm). The bacteria were then harvested and washed in 10 mL of M9 minimal medium (Nalgene). The iron levels in M9 medium were determined by inductively coupled plasma-mass spectrometry to be less than 1 nM. Following centrifugation, the bacterial pellet was resuspended in 10 mL of M9 medium and used to inoculate 50 mL of fresh M9 low-iron medium to a starting *A*_600_ of 0.04. Cultures were grown at 37 °C with shaking for 3 h before the addition of supplements (0 h) and incubated for a further 6 h. When required, antibiotics were used at the following final concentrations (μg mL^-1^): ampicillin (Ap), 100; tetracycline (Tc), 10 (for *E. coli*) and 150 (for *P. aeruginosa*); gentamicin (Gm), 250; and carbenicillin (Cb), 500.

### Gene expression analysis using quantitative real-time PCR (qRT-PCR)

To analyze gene expression, total RNA was purified from 1 mL aliquots collected at several timepoints from cultures grown under various conditions. RNA was stabilized by the addition of 250 µL of RNALater Solution (Ambion), and the samples were stored at −80 °C until further use. Total RNA was isolated from each cell pellet using the RNeasy mini spin columns according to the manufacturer’s directions (Qiagen). 6µg of total RNA was treated with RNase-free DNase I (New England Biolabs) for 3 h at 37 °C to remove contaminating chromosomal DNA and precipitated with 0.1 volume of 3 M sodium acetate (pH 5.2) and 2 volume of 100% (v/v) ethanol. RNA quantity and quality were assessed by UV absorption at 260 nm in a NanoDrop 2000c spectrophotometer (ThermoFisher Scientific). cDNA was generated using the GoScriptTM reverse transcriptase kit (Promega) from RNA (250 ng) and random primers (0.5 µg). cDNA (10 ng) was analyzed with gene-specific primers (Table S2) using the StepOnePlus realtime PCR system (Applied Biosystems) and FastStart Universal Probe Master (Roche Applied Science). The relative gene expression was calculated using the ΔΔ*Ct* method, and the cycle threshold values at each timepoint were normalized to the constitutively expressed rRNA *16s* gene. mRNA values represent the standard deviation of three independent experiments performed in triplicate.

### SDS-PAGE and Western blot analysis

Aliquots (1 ml) from PAO1 and *hasR* mutant cultures were collected at the 0-, 2-, 4-, and 6-h timepoints. Samples were harvested at 14800 rpm for 5 min at room temperature. Cell pellets were resuspended in 200 μL per 1.0 *A*_600_ of Bugbuster (Novagen). Cells were incubated at room temperature for 30 min with occasional agitation to ensure complete cell lysis, and total protein concentrations were determined using the Bio-Rad RC DC assay. Samples of the cell lysate (5 µg of total protein) in SDS-PAGE loading buffer were run on a 7.5% SDS-PAGE for HasR and PhuR detection. Following normalization between strains for differences in *A*_600_ at each timepoint, the supernatants (final volume 1-8 μL) were loaded in SDS-PAGE loading buffer. Proteins were transferred by electrophoresis to polyvinylidene difluoride membranes (Bio-Rad) for Western blot analysis. Membranes were blocked with blocking buffer (5% w/v skim milk in Tris-buffered saline (TBS) with 0.2% v/v Tween 20), washed, and probed with a 1:500 dilution of anti-HasR or anti-PhuR primary antibodies in hybridization buffer (1% w/v skim milk in TBS with 0.2% v/v Tween 20). Antibodies were obtained from Covance custom antibodies and generated from purified proteins supplied by our laboratory. Antibody sensitivity was checked against the respective purified proteins prior to use, and all experimental Western blots were run with molecular weight markers as standards. Membranes were rinsed three times in TBS with 0.2% (v/v) Tween 20 and probed with goat anti-rabbit or goat anti-mouse immunoglobin G conjugated to horseradish peroxidase (KPL) at a dilution of 1:10,000 in hybridization buffer. Proteins were visualized and enhanced by chemiluminescent detection using the SuperSignal chemiluminescence kit (Pierce) and hyperfilm ECL (Amersham Biosciences). The normalized density represents the relative abundance of each protein compared to RNA polymerase α subunit (RNApolα) as the loading control for *n =* 3 independent biological replicates. The RNApolα subunit was detected with the anti-*E. coli* antibody (BioLegend). Densitometry analysis was performed on an AlphaImager HP system using the manufacturer’s supplied AlphaView software. The HasAp supernatant samples were analyzed on the Protein Simple Wes capillary western system. No more than 4 µL of each sample was loaded with an antibody dilution of 1:25 in antibody diluent. The samples were separated by size on a 12-230 kDa capillary tray. The separation time (min), separation voltage (V), antibody diluent time (min), primary antibody time (min) and secondary antibody time (min) were 25, 375, 30, 30 and 30 respectively.

Samples were processed and quantification performed using the Compass for SW (ProteinSimple) software, where the area of each peak based on molecular weight was calculated and normalized to OD600 for (*n =* 3) independent biological replicates.

### Construction of the PAO1 hasR allelic strains

The in-frame *P. aeruginosa hasR* mutant allelic strains were constructed using two primers, *hasR upstream* and *hasR downstream* (Table S2), designed to amplify the 5’ upstream and 3’ downstream regions of the gene from the genomic DNA. The resulting Taq PCR product was ligated into pCR2.1-TOPO-TA vector (Invitrogen) for subcloning. Mutagenesis was performed by PCR using the Quikchange mutagenesis kit (Agilent Technologies) and primers specified in Table S2. The sequence of all PCR products and amino acid changes was confirmed by DNA sequencing (Eurofins MWG Operon). Following *KpnI*–*XbaI* digestion, the *hasR* mutant genes were cloned into the counter-selective suicide plasmid pEX18Tc (24). The resulting pEX18Tc*hasR* was transformed into *E. coli* S17-1-λ*pir* cells and conjugated into *P. aeruginosa* PAO1. The resulting colonies that underwent the first recombination event were screened on PIA + Tc plates at 37°C overnight. Isolated colonies were patched several times to ensure Tc^R^. To remove the pEX18Tc backbone, isolated colonies were grown in 1 mL LB shaking overnight at 210 rpm at 37°C. A 10 µL aliquot of cells was used to inoculate fresh LB cultures and was repeated twice more to induce the second recombination event. Lastly, a small dilution of cells was streaked on to a PIA + 5% sucrose plate and incubated at 37°C overnight. Several isolated colonies were patched on to a PIA + 5% sucrose and PIA + Tc plate. Only the Tc^S^ colonies were selected for sequencing to confirm complementation with the mutant allele at the original locus.

### Preparation and isolation of ^13^C-heme

Labeled [4-^13^C] δ-aminolaevulinic acid (ALA) awas purchased from Frontier Scientific. [4-^13^C] δ-ALA was used as a biosynthetic precursor to produce ^13^C-labeled heme. Expression of cytochrome *b*5 in the presence of [4-^13^C] δ-ALA induces heme biosynthesis in *E. coli* where the produced cytochrome *b*5 captures and acts as a reservoir for the synthesized heme (25). ^13^C-heme was prepared as previously reported (9, 26). Briefly, rat outer mitochondrial cytochrome *b*5 was expressed in *E. coli* BL21(DE3) cells in M9 minimal medium with the following supplements: 2 mM MgSO_4_, 100 μM CaCl_2_, 150 nM (NH_4_)_6_Mo_7_O_24_, 40 μM FeSO_4_ (acidified with 1 N HCl), 17 μM EDTA, 3 μM CuSO_4_, 2 μM Co(NO_3_)_2_, 7.6 μM ZnSO4, 9.4 μM Na_2_B4O_7_·10H2O, and 1 mM thiamine. Following lysis and centrifugation the cellular lysate was loaded onto a Q-Sepharose anion exchange column (2.5 × 10 cm) equilibrated in 50 mM Tris-HCl (pH 7.4), 5 mM EDTA, 50 mM NaCl. The column was washed with the same buffer followed by an additional wash with 50 mM Tris-HCl (pH 7.4), 5 mM EDTA, 125 mM NaCl (5 column volumes). Cytochrome *b*5 was eluted in 50 mM Tris-HCl (pH 7.4), 5 mM EDTA, 250 mM NaCl and dialyzed overnight in 50 mM Tris-HCl (pH 7.4), 5 mM EDTA, 50 mM NaCl. Heme extraction from cytochrome *b*5 was performed by the acid/butanone method as previously described (27). Heme concentrations were determined by the pyridine hemochrome assay (28). Heme stocks were prepared immediately prior to use by dissolving in 0.1 N NaOH and buffered to pH 7.4 with 1 M Tris-HCl and the final concentration was determined by pyridine hemochrome (pH 7.4). The labeling pattern of biosynthesized heme using [4-^13^C] δ-ALA is as previously reported (9).

### Extraction of BVIX isomers from P. aeruginosa supernatants

PAO1 WT and allelic strains were grown as described above and supplemented with either 1 μM ^13^C-labeled heme or HasAp bound to ^13^C-labeled heme. 50-ml cultures were shaking at 210 rpm for 3 h at 37 °C in 250-ml baffled flasks. After 3 h they were supplemented and allowed to shake for an additional 4 h. Cells were harvested by centrifugation in 50 mL conical tubes at 6000 rpm for 20 min at 4°C. The BVIX isomers were extracted as previously described (16). Briefly, the supernatant was collected and filtered by a Nalgene vacuum filtration unit with a 0.22 µm PVDF membrane. Supernatants were acidified to pH ∼ 2.5 with 10% trifluoroacetic acid (TFA), supplemented with 10 nM (final concentration in 45 mL) with Di-Methyl ester BVIXα and loaded over a C_18_ Sep-Pak column purchased from Waters. The column is prepared by flushing with 2 mL of Acetonitrile (ACN), H_2_O, 0.1% TFA in H_2_O and methanol:0.1% TFA (10:90). After sample application wash with 4 mL 0.1% TFA, 4mL ACN:0.1% TFA (20:80), 2 mL Methanol:0.1% TFA (50:50). Elute with 1 mL Methanol. Purified BVIX isomers were speed vacuumed dry and stored for up to 1 week at −80 °C prior to LC-MS/MS analysis.

### LC-MS/MS Analysis of BVIX Isomers

Samples were resuspended in 10 μL DMSO and diluted to 100 μL with mobile phase ACN (50:50, v/v) and centrifuged at 14000 rpm for 5 minutes at room temperature to remove particulates. The BVIX isomers (2 μL) were separated and analyzed by LC-tandem Mass Spectrometry (MS/MS) (Waters TQ-XS triple quadrupole mass spectrometer with AQUITY H-Class UPLC) as previously described with slight modification (16). BVIX isomers were separated on an Ascentis RP-amide 2.7 µm C18 column (10 cm × 2.1 mm) at a flow rate of 0.4 mL/min. The mobile phase consisted of A: H_2_0:0.1% formic acid and B: ACN:0.1% Formic acid. The initial gradient is 64% A and 36% B. After 5 min A:55% - B%45, 8 min A:40% - B:60%, 8.5 min A:5% - B:95% and 10 min A:64% - B:36%. Fragmentation patterns of the precursor ions where detected at 583.21 (^12^C-BVIX) and 591.21 (^13^C-BVIX) using multiple reaction monitoring (MRM). The source temperature was set to 150°C, the capillary voltage to 3.60 kV and the cone voltage to 43V. The column was kept at 30°C during separation. The precursor ion used in MRM for ^12^C-BVIXα, -β, and -δ isomers is 297.1, 343.1, and 402.2 with a collision energy of 38, 36 and 30 V respectively. The product ion used in MRM for ^13^C-BVIXα, -β, and -δ isomers is 301.1, 347.1 and 408.2 with a collision energy of 34 V for each.

### ICP-MS analysis of intracellular iron content

Aliquots (2 ml) were removed from cultures grown as described for the heme uptake studies at 2, 5 and 7 h, pelleted and washed in fresh M9 media. The pellets were dissolved in trace-metal-free ultrapure 20% HNO_3_ and boiled overnight at 100°C. Samples were further diluted with ultrapure water to a final concentration of 2% HNO_3_ and subjected to ICP-MS on Agilent 7700 ICP-MS (*Agilent Technologies*). ICP-MS runs were calibrated with high-purity iron standard solution (*Sigma-Aldrich*) and raw ICP-MS data (ppb) were corrected for drift using values for scandium and germanium (*CPI international*) as internal standards and added to samples during processing. Corrected values were then normalized to culture density as determined by the absorbance at 600 nm. Experimental values and reported standard deviation were the average of three biological replicates.

## SUPPLEMENTAL MATERIAL

Supplemental material for this article can be found at Table S1 Table S2 Fig S1 Fig S2 Fig S3

## ACKNOWLEDGMENTS

The authors thank the University of Maryland, School of Pharmacy, Mass Spectrometry Center including the Center Director Maureen Kane and Stephanie Shiffka for their technical expertise. ATD would also like to thank Susana Mouriño for helpful discussion and reading of the manuscript.

## AUTHOR CONTRIBUTIONS

ATD performed the experiments and analyzed the data. ATD and AW wrote and edited manuscript.

## FOOTNOTES

This work was supported by National Institutes of Health Grant R01 AI134886 (to A. W.) ATD was funded by National Institutes of Health Ruth L. Kirschstein (NRSA) Individual Predoctoral Fellowship GM126860. ATD also wishes to acknowledge previous support from a Chemistry/Biology Interface Training Program (NIGMS/NIH T32GM066706).

